# Roles of PknB and CslA in cell wall morphogenesis of *Streptomyces*

**DOI:** 10.1101/2024.08.02.606377

**Authors:** Marta Derkacz, Andrew Watson, Akshada Gajbhiye, Michał Tracz, Dagmara Jakimowicz, Matthias Trost, Jeff Errington, Bernhard Kepplinger

**Affiliations:** University of Wroclaw, Department of Molecular Microbiology, ul. F. Joliot-Curie 14a, 50-383 Wrocław, PL; University of Newcastle, Centre for Bacterial Cell Biology, Baddiley-Clark Building, Newcastle upon Tyne NE2 4AX, UK; Newcastle University Biosciences Institute, Faculty of Medical Sciences, Newcastle upon Tyne, NE2 4HH, UK; University of Wroclaw, Mass Spectrometry Laboratory, ul. F. Joliot-Curie 14a, 50-383 Wrocław, PL; The University of Sydney, Faculty of Medicine and Health, Sydney, NSW 2006, AU

**Keywords:** DivIVA, PknB, CslA, Tip Organising Centre (TIPOC), polarisome, cell morphogenesis, cell wall, phosphorylation, *Streptomyces*

## Abstract

The bacterial cell wall is essential for maintaining cellular integrity and defining the mode of growth, with different species adopting distinct strategies for cell wall synthesis and remodelling. *Streptomyces* are filamentous bacteria predominantly found in soil and renowned for producing specialised metabolites, including antibiotics. They grow through tip extension and branching hyphal filaments, forming a multicellular mycelium. New branches are established by forming a new growth zone on the lateral cell wall. Proteins involved in this process are organised into complexes called polarisomes, with DivIVA being the most well-characterised component. To investigate the tip growth requirements in *Streptomyces albus* we developed a genetic screen utilising toxic DivIVA overproduction and searched for suppressors of its lethality, reasoning that such suppressors would likely encode components functionally linked to DivIVA or the tip growth machinery. Among the identified genes was *pknB*, encoding a serine/threonine protein kinase implicated in the regulation of cell growth and morphogenesis. We confirmed that deletion of *pknB* restored the growth phenotype of *S. albus* following DivIVA overproduction. The phosphoproteome analysis revealed that the absence of PknB alters the phosphorylation state of CslA, a cellulose synthase-like protein. We demonstrate that a phosphoablative mutant of CslA impairs β-glucan synthesis and causes hypersensitivity to lysozyme. Overproduction of CslA restored colony growth defects arising from DivIVA-induced hyperbranching, without however suppressing the hyperbranching phenotype.

These findings collectively identify PknB-dependent phosphorylation of CslA as a central regulatory point in *Streptomyces* cell envelope construction, revealing how modulation of β-glucan synthesis can mitigate the cellular consequences of DivIVA dysregulation.

## Introduction

Peptidoglycan, a major target for antibiotics, is an essential component of bacterial cell walls, providing structural integrity and rigidity and enabling bacteria to maintain their shape and withstand osmotic pressure challenges (Liu and Breukink, 2016; Garde *et al*., 2021). Proteobacteria (e.g., *Escherichia coli*) and Firmicutes (e.g., *Bacillus subtilis*) are among the best-studied models for cell morphogenesis. These rod-shaped bacteria typically exhibit lateral insertion of new cell wall material during elongation (Egan *et al*., 2020). In contrast, Actinobacteria (e.g., *Streptomyces* and *Mycobacteria*) mainly insert new cell wall material at the tips during exponential growth (Flärdh, 2003a; Kuru *et al*., 2014; Zambri *et al*., 2025). Actinobacteria can be unicellular, such as Mycobacteria or Corynebacteria, or multicellular (e.g., *Streptomyces*), the latter forming a complex mycelial network that supports substrate colonisation (Chater, 2006). This polar growth mode is hypothesised to confer ecological advantages, such as efficient nutrient foraging and competitive dominance in soil environments.

Cell wall biosynthetic proteins driving tip growth are organised locally into polarisome complexes (Hempel *et al*., 2008). Within the polarisome, DivIVA—a coiled-coil protein—plays a pivotal role and is essential for *Streptomyces* growth (Flärdh, 2003b). Overproduction of DivIVA results in swollen tips and hyperbranching, likely due to the misregulation of the cell wall biosynthetic machinery (Flärdh, 2003b). New DivIVA complexes appear on the lateral wall preceding the formation of the new branches (Flärdh, 2003b; Hempel *et al*., 2008; Richards *et al*., 2012). Two additional scaffolding proteins, Scy and FilP, are believed to sequester DivIVA, thereby facilitating the establishment of new growth zones and influencing the size and position of DivIVA complexes, respectively (Holmes *et al*., 2013; Fröjd and Flärdh, 2019). The precise mechanism for branch initiation remains however elusive (Hempel *et al*., 2008).

DivIVA phosphorylation has been reported in a wide range of species (Kang *et al*., 2005; Beilharz *et al*., 2012; Elsholz *et al*., 2012). *Streptomyces coelicolor* encodes at least 34 Ser/Thr kinases, and DivIVA is modified by the non-essential Ser/Thr kinase AfsK (Hempel *et al*., 2012). Mass spectrometry has identified AfsK phosphorylation sites on DivIVA as occurring on the non-essential C-terminal protein at residues T304, S309, S338, S344, and S355 in *S. coelicolor* (Saalbach *et al*., 2013).The phosphatase SppA appears to function as a counterpart in this process (Passot *et al*., 2022). Notably, AfsK overexpression induces disassembly of the apical (mother) polarisome, leading to the formation of multiple daughter polarisomes, followed by the establishment of multiple hyphal branches (Hempel *et al*., 2008).

In contrast, in Mycobacteria, the DivIVA homologue Wag31 is phosphorylated by the essential kinases PknA and PknB (Jani *et al*., 2010). These genes reside within the division and cell wall gene cluster that encodes key components of the bacterial cell division and peptidoglycan biosynthesis machinery (Molle and Kremer, 2010). It typically includes *fts* genes for Z-ring assembly, *mur* genes for peptidoglycan precursor synthesis, *rod*-*pbpA* genes for the cell wall, and regulatory serine/threonine kinases *pknA* and *pknB*. Interestingly, the elimination of the homologous kinase PknB in *Streptomyces* has no detrimental effect, whereas the kinase PknA is missing in most *Streptomyces spp.* (Jones *et al*., 2011).

In this study, we investigated the consequences of DivIVA overproduction in *Streptomyces albus* to uncover the factors that are involved tip growth. This study revealed that inactivation of the kinase PknB unexpectedly alleviated defects resulting from DivIVA imbalance. Subsequent analysis identified CslA as a key downstream target of PknB, a cellulose synthase–like protein that synthesises β-glucan and contributes to cell wall biogenesis (Xu *et al*., 2008). We further demonstrate that CslA phosphorylation state modulates β-glucan-synthesis and, consequently, the cell’s ability to stabilise growth zones. These findings suggest a phosphorylation-dependent regulatory mechanism linking PknB activity to cell wall assembly at hyphal tips.

## Materials and methods

### Bacterial strains and culture conditions

All *S. albus* strains used in this study are listed in Table S1. *S. albus* spores were collected using standard protocols (Kieser T *et al*., 2000) from cultures grown on Soya Flour Mannitol (SFM) agar plates supplemented with antibiotics (if required) to a final concentration of apramycin (50 µg/mL) and hygromycin (50 µg/mL). Spore concentration was determined by serial dilution on SFM agar plates. Spores were maintained at–80 °C in 25% glycerol aliquots for single-use. Tryptic Soy Broth (TSB) and 5% TSB were used for morphology experiments. Liquid cultures were performed in 250 mL Erlenmeyer flasks containing metal spirals at 30 °C and 220 rpm for agitation. Solid cultures were grown on 20 mL of 2% or 5% TSA agar plates at 30 °C. *E. coli* strains (listed in Table S2) were grown at 37 °C on LB supplemented with antibiotics (if required) to a final concentration of apramycin (50 µg/mL), kanamycin (50 µg/mL), chloramphenicol (34 µg/mL), or on 2YT supplemented with hygromycin (50 µg/mL).

### Construction of plasmids and strains

All plasmids and oligonucleotides used in this study are listed in Table S3 and Table S4, respectively. Details of the plasmid construction are provided in the Supplementary Methods, specifically in the section on comprehensive plasmid design and assembly. Plasmids were transformed into *E. coli* DH5α and subsequently transferred to *Streptomyces* by conjugation with *E. coli* ET12567/pUZ8002 strain or protoplast transformation using standard protocols (Kieser T *et al*., 2000). For integrative plasmids, *Streptomyces* strains were grown in the presence of the appropriate antibiotics. CRISPR/Cas9 plasmids were cured by incubation at 42 °C without antibiotics. Mutants lacking apramycin resistance were sequenced by Microsynth AG using PCR-amplified fragments of modified genomic DNA (with the exception of the 65α, 65α attBϕC31:: pIJ6902 (empty plasmid), and 65α attBϕC31::pBK288 (CslA under *tipA* inducible promoter) strains, which were verified by Illumina sequencing).

### Selection of spontaneous suppressor mutants and bioinformatics analysis

Spores of the DivIVA overproduction strain (approximately 10^6^ spores per plate) were spread onto 5% TSA plates supplemented with 500 ng/mL of inducer (anhydrotetracycline). Following 5 days of incubation, several colonies were observed and subsequently restreaked onto fresh 5% TSA containing 500 ng/mL of the inducer (anhydrotetracycline). The parent DivIVA-overproducing and wild-type strains served as controls.

DNA was isolated from the strains using a modified version of the “salting-out” method (Pospiech and Neumann, 1995; Watson *et al*., 2022). Libraries for Illumina sequencing were prepared using the Illumina DNA prep kit, Nextera XT kit, and Illumina Nextera XT primer Index Kit. Sequencing was performed by Edinburgh Genomics using MiSeq v2 (250 PE). Raw sequences are available in the NCBI Sequence Read Archive (accession number: PRJNA1099611).

The quality of all Illumina reads was assessed using fastQC (version 0.11.9), and the reads were trimmed using trimmomatic (version 0.39). Bowtie2 (version 2.3.5.1) was used to align the trimmed Illumina reads to the *Streptomyces albus* J1074 genome (NC_020990.1) using the very sensitive preset mode for mapping paired-end reads. Variants were called using bcftools (version 1.14) in the consensus calling mode (-c). Only predicted variants with a QUAL score of ≥ 15 were retained. Bedtools subtract (version 2.27.1) was used to filter out any variants identified in potential suppressor strains that were already present in the lab *S. albus* J1074 parent strain compared to the reference sequence (NC_020990.1) and thus were accumulated independently from this experiment. The potential impact of all variants on protein-coding sequences was assessed using snpEff (version 5.0e) (Cingolani *et al*., 2012). The final table was derived by manually inspecting shortlisted mutations.

### Microscopy

For phenotypic analysis, 10^7^ colony-forming units (CFU) of *S. albus* J1074 strains were inoculated into 20 mL TSB or 5% TSB, with or without 250 ng/mL anhydrotetracycline (ATc), and cultured for 20 h.

For β-glucan staining, 2·10^6^ CFU (estimated by OD600 measurement) of *S. albus* J1074 strains from freshly grown SFM agar plates were inoculated in 20 ml TSB and cultured for 20 h. Cells were centrifuged and washed with modified PEM buffer (0.4 M PIPES, 20 mM EDTA, 20 mM MgCl2; pH = 6.9 (KOH)), stained with lectin dye (Sigma-Aldrich L3892) for 5 min at room temperature, washed, and resuspended in modified PEM buffer (Hasek, 2006).

Cells were centrifuged as needed, loaded onto a 1.2% agarose slide prior to imaging, and examined using a Leica DM6 B fluorescence microscope equipped with a 100× objective and a DFC7000 GT camera. Images were analysed using the ImageJ software.

To determine the β-glucan content (Fig. 3), the mean fluorescence signal was measured at each tip (within a 20-pixel diameter circle). A minimum of 574 data points were collected for each strain tested. Data were analysed and visualised using R software, including the ggplot2 package (Ehrlinger, 2016). Statistical analysis was performed using analysis of variance (ANOVA), followed by Tukey’s HSD post-hoc test.

### Protein extraction for mass spectrometry

For mass spectrometry, 10^7^ CFU of WT or Δ*pknB* were inoculated in 50 mL of TSB or 5% TSB and cultured for 20 h. Four biological replicates were performed under each condition. Mycelia were harvested by centrifugation (5 min, 4 °C, 3000 × g), washed twice with phosphate-buffered saline (PBS), and resuspended in 8 M urea buffer supplemented with 50 mM Tris, 1 mM TCEP (Roth), a protease inhibitor (Thermo Scientific), and a phosphatase inhibitor consisting of sodium molybdate (115 mM), sodium orthovanadate (100 mM), sodium tartrate dihydrate (400 mM), glycerophosphate (500 mM), and sodium fluoride (100 mM). Five millilitres of buffer was used per gram of biomass. Cell disruption was achieved via sonication (2 × 20 s, 4 °C, 20 kHz), followed by centrifugation (15 min, 4 °C, 3000 × g), and the resulting supernatant was stored at–80 °C prior to subsequent steps. A detailed protocol for further sample preparation and instrumental analysis is provided in the Supplementary Information.

### Lysozyme sensitivity and survival assays

Sensitivity to lysozyme was assessed by inoculating 5 µL of PBS spore suspensions and 1:10 dilutions onto TSA plates supplemented with lysozyme at a final concentration of 25 µg/mL, compared to a no-addition control (Fig. 3A). Cultures were incubated for 3 days. Survival assays were performed similarly using TSA and 5% TSA, both supplemented or not with anhydrotetracycline at 500 ng/mL. To determine whether osmotic stress conditions induced by 5% TSA affected the lethality of the inducible DivIVA production strain (strain 65α), 0.5 M sucrose was added to the 5% TSA.

### DivIVA-mScarlet production analysis

For western blot analysis, 2 × 10^7^ CFU of *Streptomyces* strains were inoculated in 50 mL of TSB or 5% TSB, in the presence or absence of ATc (250 ng/mL), and cultured for 24 h. Up to 50 mL of cultured mycelia were harvested by centrifugation (5 min, 4 °C, 3000 × g), washed twice with PBS, and resuspended in 1 mL PBS supplemented with a protease inhibitor (Thermo Fisher Scientific) and 1 mM DTT. Cell disruption was achieved via sonication (2 × 20 s, 4 °C, 20 kHz), followed by mixing with 6x loading buffer and denaturation (15 min, 95 °C). Total protein concentration was normalised and assessed using SDS-PAGE containing 2.5% 2, 2, 2-trichloroethanol (Thermo Scientific). Proteins were visualised using a stain-free filter on a ChemiDoc (Bio-Rad). Signal intensity was measured using ImageJ software, and the samples were adjusted accordingly. Proteins were transferred to a nitrocellulose membrane, which was blocked with a blocking solution (5% powdered milk in PBST [PBS with 0.1% Tween-20]), then incubated with either rabbit anti-mCherry polyclonal antibody (diluted 1:1000 in fresh blocking solution) or anti-DivIVASC antiserum (diluted 1:5000 (gift from Prof, Klas Flärdh, Lund University) (Wang *et al*., 2009). The membrane was washed thrice with PBST and incubated with goat anti-rabbit IgG cross-linked to horseradish peroxidase (Santa Cruz Biotechnology; diluted 1:5000 in fresh blocking solution), followed by three washes with PBST. The blot was incubated with SuperSignal ™ West Pico PLUS Chemiluminescence Substrate (ThermoScientific) and visualised using a ChemiDoc (BioRad).

### Growth Assay with Induced *cslA* Mutant

Overnight cultures were harvested by centrifugation and washed thrice with phosphate-buffered saline (PBS). The optical density (OD) of the washed cells was adjusted to 0.1, and aliquots were plated onto 5% tryptic soy agar (TSA) plates supplemented with either 500 ng/mL of the DivIVA inducer and/or 0.5 µg/mL of thiostrepton.

## Results

### Identification of conditions for lethality of a DivIVA overexpression construct

The scaffolding protein DivIVA is localised at the tip of the *Streptomyces* hyphae and is essential for regulating normal apical growth (Flärdh, 2003b; Hempel *et al*., 2008). Previous studies have demonstrated that overproduction of DivIVA leads to hyphal tip swelling and overbranching (Hempel *et al*., 2008). We hypothesised that lethal overproduction of DivIVA could be suppressed by mutations in genes encoding proteins involved in tip growth and cell wall remodelling. Initially, we constructed a strain carrying a *divIVA-mScarlet* fusion under the control of an anhydrotetracycline (ATc)-inducible promoter located in the phiBT1 locus. Although induction of this fusion protein was non-lethal, during the development of additional derivatives, we serendipitously created a strain, designated 65α, harbouring two inducible copies of *divIVA-mScarlet*, one in the native locus and the other at the phiBT1 locus (Fig. 1A). This strain was able to grow in the absence of the inducer, presumably because of leaky expression from the promoter (Fig. 1B). However, upon induction, a strong fluorescence signal was observed at the hyphal tips, accompanied by swelling and hyperbranching (Fig. 1C), aligning with previously documented observations from a DivIVA overproduction strain of *S. coelicolor* (Flärdh, 2003b; Hempel *et al*., 2008). After testing a range of growth conditions, we found that under hypoosmotic stress conditions (1/20 reduced concentration of TSA medium “5% TSA”), the growth of strain 65α was abolished in the presence of the inducer, while it grew similarly to the wild type in its absence (Fig. 1B), suggesting cell wall damage leading to osmotic lysis (rescued by 0.5 M sucrose; Fig. S1).

**Figure 1.**
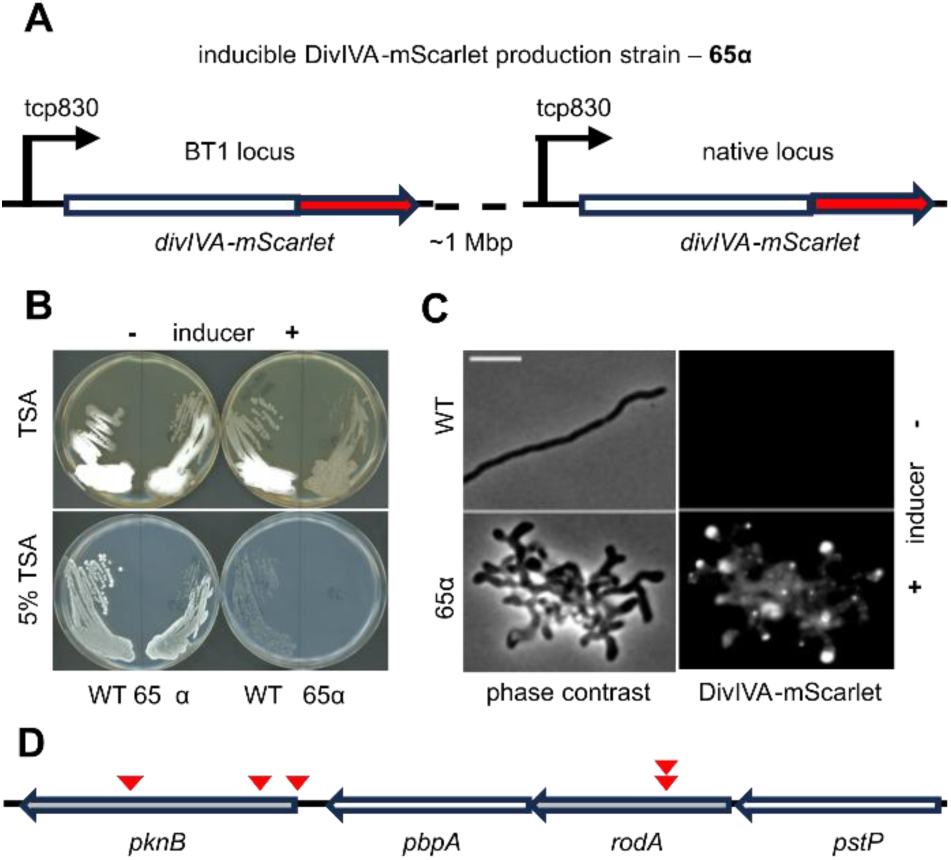
Genetic configuration and phenotype of DivIVA overproducing strain 65α (A) Pictorial representation of the genetic makeup of the *S. albus* J1074 mutant, called 65α, containing two copies of a *divIVA-mScarlet* fusion under the control of the ATc inducible promoter tcp830. (B) Comparison of the growth behaviour of the wild-type *S. albus* J1074 and the inducible mutant strain 65α on TSA and 5% TSA in the presence or absence of the inducer (500 ng/mL). (C) Microscopic phenotypes of *S. albus* J1074 wild-type strain and 65α mutant grown in TSB in the presence of the inducer (250 ng/mL). Fluorescent phase-contrast microscopy demonstrated DivIVA-mScarlet localisation. Scale bar - 5 µm. (D) Genetic organisation of the gene cluster containing *pknB* and *rodA* in *S. albus* J1074 (3,450,591-3,457,165 on NC_020990; ∼1Mbp from BT1 locus and ∼2Mbp from *divIVA* locus). The red arrows indicate the locations of the mutations described in Table 1.

**Table 1:** Genes recurrently mutated across independent suppressor mutants.

| Gene Name | Protein names | Mutant number<sup>a</sup> | Amino acid change<sup>b</sup> |
| --- | --- | --- | --- |
| *XNR_RS08380* | prolyl serine peptidase | 2 | M34:P63fs; M43:P63fs |
| *XNR_RS15060* | PknB | 3 | M25: M1fs; M40: A396D; M43: S98fs |
| *XNR_RS15070* | RodA | 2 | M33: L162del; M34: L162del |

<sup>a</sup> Mutant number refers to the frequency at which a mutation in a gene was recovered in a different mutant strain. The amino acid change indicates the result of a mutation in the corresponding mutant strain starting with M.
<sup>b</sup> fs – frame shift; del – deletion

### A suppressor screen for mutations bypassing *divIVA* overexpression lethality

To identify genes potentially influencing hyphal growth, we plated *divIVA* overexpression strain 65α under the lethal conditions described above and picked the rare colonies that emerged. Of the 43 colonies from three different starting cultures, 13 mutant strains displayed robust growth under 65α lethal conditions.

Because we had two separate copies of the *divIVA* construct, we expected the suppressor mutations to lie in genes other than *divIVA*. Whole-genome sequencing followed by bioinformatic analysis of the 13 mutants revealed apparent mutations in 122 coding regions (File S1). Table 1 lists the genes in which more than one mutation was identified. Of these, XNR_RS15060 and XNR_RS15070 were particularly interesting. They are located almost adjacent to each other in the division and cell wall (DCW) cluster, and their identities are apparent from the conservation of the gene cluster relative to other actinobacteria. We annotated these as *pknB* (XNR_RS15060) and *rodA* (XNR_RS15070) based on their homology to the equivalent operon in *S. coelicolor* (Fig. 1D). In *pknB*, we identified two distinct frameshift mutations: one affecting the start codon and the other located within the kinase domain near the N-terminus. An additional variant of this gene consisted of a single nucleotide substitution within the PASTA domain (Fig. S2).

PknB kinase is an essential protein in *Mycobacterium tuberculosis* (Chawla *et al*., 2014) and *Corynebacterium glutamicum* (Fiuza *et al*., 2008), and conditional *pknB* mutations affect the maintenance of cell shape (Arora *et al*., 2018). PknB also phosphorylates DivIVA in Mycobacteria, but not in *Streptomyces* (Kang *et al*., 2005; Hempel *et al*., 2012). The exact role of PknB in *Streptomyces* has not been elucidated, as deletion of *pknB*, in contrast to other bacteria, has no deleterious effect, at least in wild-type cells (Jones *et al*., 2011). The network of interactions described above, involving various proteins involved in cell morphogenesis, prompted us to focus on the role of PknB.

### Deletion of *pknB* rescues *Streptomyces* from lethal overexpression of *divIVA*

To test whether the mutations identified above were responsible for the suppression of the *divIVA* overexpression phenotype, we attempted to delete *pknB* in wild-type and *divIVA* overexpression (65α) strains. As expected, deletion of *pknB* did not result in any evident phenotypes in the wild-type *S. albus* strain, consistent with previously reported results (Jones *et al*., 2011). However, the 65α background deletion of *pknB* enabled growth under otherwise lethal conditions (5% TSA + ATc) (Fig. 2A).

**Figure 2.**
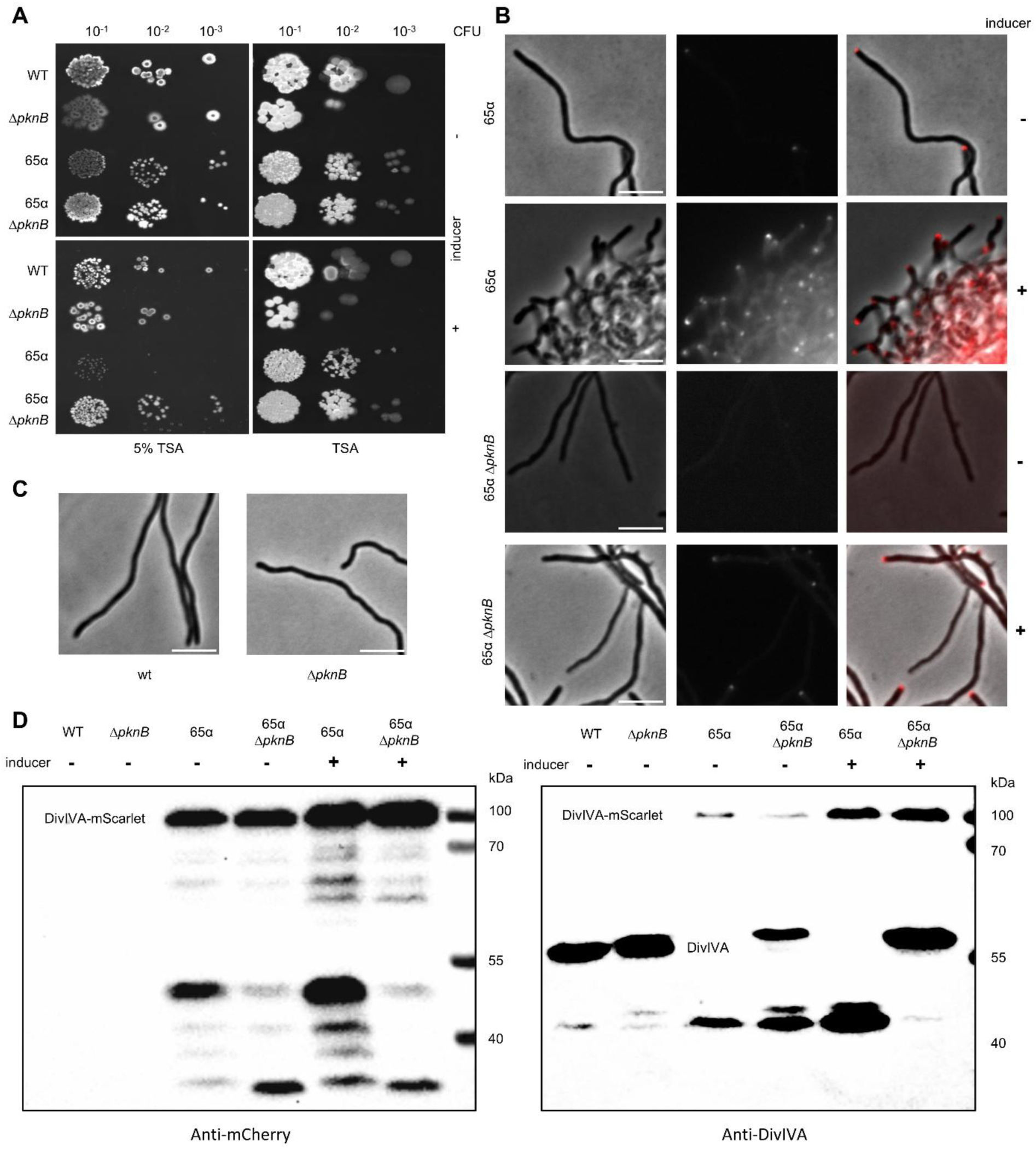
Verification of *pknB* deletion as a suppressor of lethal DivIVA overproduction. (A) Survival assay of *S. albus* J1074 strains grown on 5% TSA or TSA in the presence or absence of the inducer - ATc 500 ng/ml. B) Phenotypes of *S. albus* J1074 strains grown in TSB in the presence or absence of the inducer ATc (100 ng ml⁻¹). Fluorescence and phase-contrast microscopy revealed DivIVA–mScarlet production. Fluorescence backgrounds and intensity thresholds were adjusted identically across samples to allow for direct comparison. In the overlay panel, fluorescence levels were adjusted for improved visualisation. Scale bar, 5 µm. (C) Phase-contrast microscopy of *S. albus* J1074 and *S. albus* J1074 Δ*pknB* grown in TSB. (D)Western blot analysis of DivIVA-mScarlet (theoretical mass ∼68 kDa) production by *S. albus* J1074 strains grown in TSB, in the presence or absence of the inducer - ATc 250 ng/ml. Left Detection was performed using anti-mCherry polyclonal antiserum. Right Detection was performed using anti-DivIVASC antiserum. The molecular weight (MW) values to the right indicate the positions of the marker proteins.

Microscopic imaging (Fig. 2B-C, quantification Fig. S3) revealed that the 65α strain had an increased frequency of branching compared with the wild type, even in the absence of the inducer, while in the presence of ATc, it exhibited hyperbranching, bulging, and an accumulation of mScarlet at the hyphal tips. In contrast, the 65α Δ*pknB* strain exhibited reduced branching and a diminished fluorescent signal compared with the 65α parent. Indeed, 65α Δ*pknB* mycelia closely resembled those of the wild-type strain, even in the presence of ATc. Although we could still detect a fluorescent signal from mScarlet, the fluorescence was notably reduced compared to that of 65α. In the absence of the inducer, most hyphae (67%) exhibited a diffuse cytoplasmic distribution of the mScarlet signal. Upon induction, this proportion decreased to approximately 33%, with the remaining hyphae displaying a clear accumulation of fluorescence at the tips of the hyphae.

Sanger sequencing of both *divIVA-mScarlet* fusion genes did not reveal any mutations. Presumably, the lowered signal at the hyphal tips could be a result of DivIVA-mScarlet being distributed more widely throughout the hyphae or diminished protein levels due to decreased synthesis or increased degradation.

To compare the total DivIVA protein levels between strains, we analysed DivIVA–mScarlet abundance in 65α and 65α *ΔpknB* by Western blotting using anti-mScarlet- and anti-DivIVA- antibodies (Fig. 2D; loading control in Fig. S4). Native DivIVA (predicted 41.1 kDa) migrated at approximately 60 kDa in both wild-type and Δ*pknB* backgrounds. Its abundance was not significantly altered in the absence of *pknB*. The upward shift in the apparent molecular weight likely reflects dimerisation or altered electrophoretic mobility. In the 65α background, the DivIVA–mScarlet fusion protein (predicted 68 kDa) migrated near 100 kDa. Upon ATc induction, the intensity of the full-length fusion increased strongly when detected by both the anti-DivIVA and anti-mScarlet antibodies, confirming that induction elevated total DivIVA–mScarlet levels. Both *65α* and *65α ΔpknB* strains displayed additional faster-migrating- species. These fragments were detected by anti-DivIVA but not by anti-mScarlet antibody, indicating that they represent N-terminal- DivIVA cleavage products lacking the fluorescent tag. Conversely, the anti-mScarlet- blot revealed smaller mScarlet-containing- fragments not recognised by anti-DivIVA-, consistent with C-terminal- cleavage events. ATc induction increased the abundance of these fragments in *65α*, consistent with the higher overall expression of the fusion protein.

Loss of *pknB* did not reduce the amount of full-length- DivIVA–mScarlet; however, it markedly altered the pattern of degradation products. The 65α *ΔpknB* strain accumulated several distinct DivIVA-sized fragments, suggesting that *pknB* influences the stability or processing of the DivIVA–mScarlet fusion protein indirectly. This raises the possibility that PknB-dependent signalling influences the proteolytic turnover of DivIVA, potentially serving as a mechanism to control DivIVA levels under conditions of overexpression.

### Deletion of *pknB* causes hyperphosphorylation of CslA

The kinase PknB has been reported to be involved in various processes across various organisms, including cell division, peptidoglycan synthesis, protein synthesis, and stress responses (Jones *et al*., 2011; Dworkin, 2015; Richard-Greenblatt and Av-Gay, 2017). To better understand its role in *S. albus,* we analysed the phosphoproteomes of wild-type and *ΔpknB S. albus* strains, with particular focus on proteins implicated in polar growth. The results identified 27 phosphorylated proteins, of which three were significantly hypophosphorylated in the *pknB* mutant, while one was significantly hyperphosphorylated (Table 2 and File S2; proteomics data are available in File S3). Among the expected candidates, we anticipated changes in DivIVA phosphorylation, as well as in other regulators such as the kinase AfsK or the phosphatase SppA, which is known to dephosphorylate DivIVA (Hempel *et al*., 2012; Passot *et al*., 2022). Although SppA phosphorylation was indeed altered in the Δ*pknB* background, no corresponding changes in DivIVA phosphorylation were detected in our experimental setup, and this pathway was therefore not investigated further. Our attention was drawn to CslA, a synthase previously associated with polar growth, whose phosphorylation was hyperphosphorylated on residue T17 in the Δ*pknB* background, making it a compelling candidate for mediating PknB’s effects on hyphal development. This protein localises at hyphal tips and is responsible for the production of a beta-glucan-containing polysaccharide that is thought to provide additional physical protection, potentially required for the continuous remodelling of the growing cell wall (Xu *et al*., 2008). Loss of this polymer impairs the morphological development and stability of the cell wall (Xu *et al*., 2008; Zhong *et al*., 2022). CslA forms a protein complex with the copper radical oxidase GlxA, and it is a known interaction partner of DivIVA in *S. coelicolor* (Xu *et al*., 2008).

**Table 2.** Significantly different protein phosphorylation patterns of *S. albus* J1074 WT and Δ*pknB* strains grown in TSB medium.

| locus_tag | protein name | phosphorylation in $\Delta pknB$ vs WT |
| --- | --- | --- |
| XNR_RS17030 | FolK | Hypophosphorylated ↓ |
| XNR_RS11880 | GroL | Hypophosphorylated ↓ |
| XNR_RS08650 | CslA | Hyperphosphorylated ↑ |
| XNR_RS14580 | SppA | Hypophosphorylated ↓ |

To evaluate the effect of phosphorylation on the function of CslA, we constructed a strain producing phosphoablative CslA (*cslA*T17A) and a deletion mutant. We confirmed the production of the phosphoablative ClsA using a proteomics experiment (Fig S5). Previous studies have shown that deletion of *cslA* increases sensitivity to lysozyme (Zhong *et al*., 2022); we therefore tested whether preventing phosphorylation at T17 would similarly affect cell wall integrity. Both *cslA* mutants displayed hypersensitivity to lysozyme, suggesting that phosphorylation is important for CslA activity (Fig. 3A). To determine whether the reciprocal change — hyperphosphorylation of CslA in the Δ*pknB* background — had the opposite effect, we assessed lysozyme sensitivity of the *pknB* deletion strain. This strain responded to lysozyme similarly to the wild type, indicating that hyperphosphorylation of CslA alone is not sufficient to further enhance cell wall protection beyond wild-type levels. Next, we used a lectin dye to stain β-glucans (Fig. 3B-C) and found that the signal at the tips of hyphae was completely abolished in the *cslA* knockout strain, similar to previously reported data on a *cslA* deletion strain (Xu *et al*., 2008; Zhong *et al*., 2022). Again, the strain producing phosphoablative variant of CslA (*cslA*(T17A)) behaved similarly to the deletion strain when stained for β-glucans, indicating that phosphorylation is likely required for the β-glucan synthetic activity of CslA. Overall, these data indicate that phosphorylation plays an important role in β-glucan production, although the role of PknB in this process is likely to be indirect.

**Figure 3.**
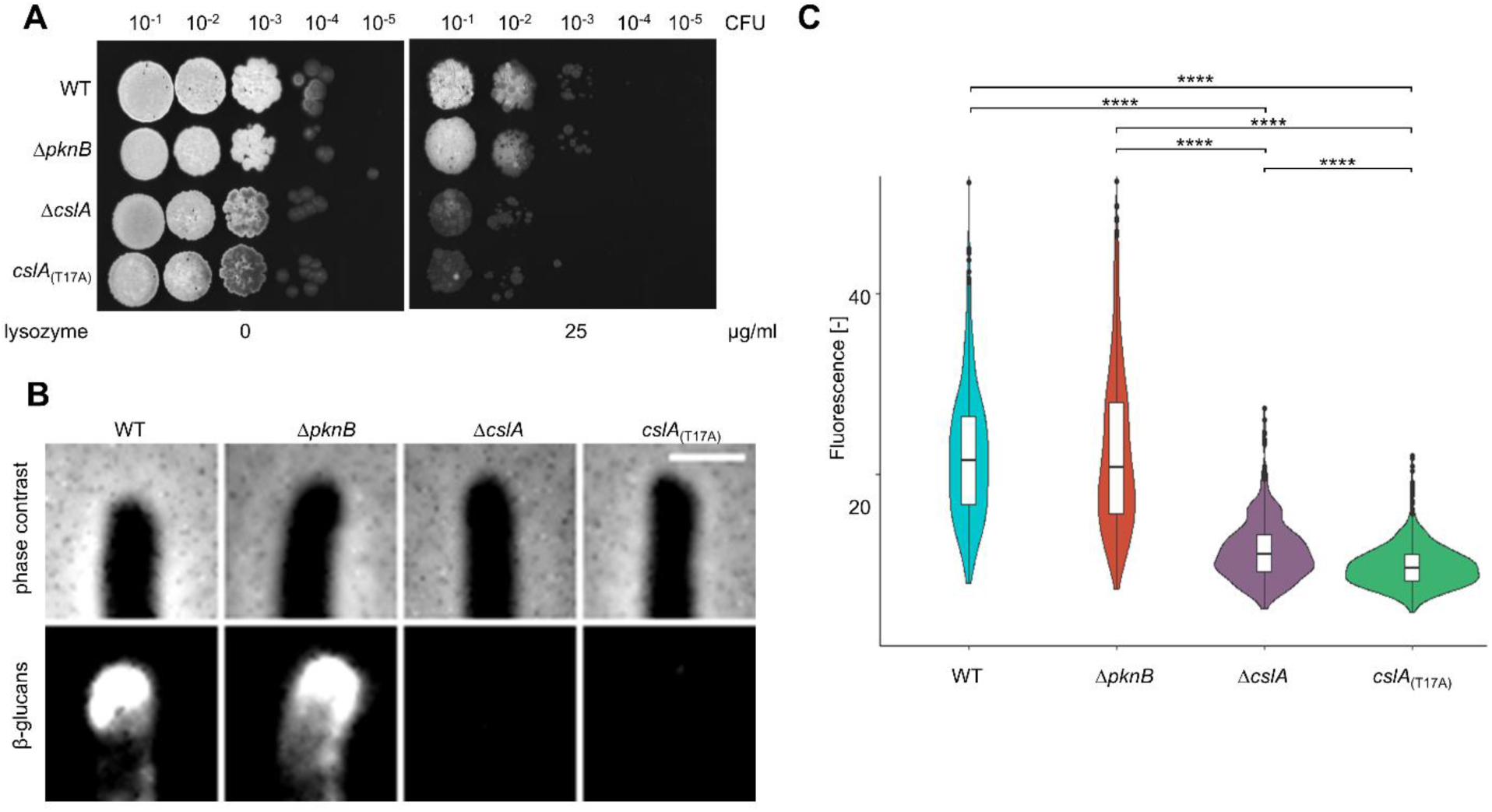
Impact of mutations in *cslA* on glucan production (A) Lysozyme sensitivity of *S. albus* J1074 strains grown on TSA. (B) Phenotypes of *S. albus* J1074 strains grown on TSA. Cells were stained with lectin dye to visualise β-glucans. Scale bar - 1 µm. (C) Quantification of mean β-glucan production at the tip by *S. albus* J1074 strains grown in TSB. The width of each plot indicates the number of collected data points for a specific fluorescence. Significant differences, according to ANOVA followed by Tukey’s HSD post-hoc test, are marked with **** (p≤0,0001).

### Overexpression of CslA rescues cells from DivIVA overproduction lethality

The results described above, showing that phosphorylation of CslA is likely required for the β-glucan synthetic activity, suggest that the deletion of *pknB* might increase the activity of CslA, via increased phosphorylation albeit indirectly. If so, *cslA* overexpression might also rescue cells from *divIVA-mScarlet* overexpression toxicity. To test this, a plasmid carrying a thiostrepton-inducible copy of *cslA* was introduced into the 65α background, with an empty plasmid as a control. In the absence of DivIVA overproduction, both strains grew well on 5% TSA. However, even without induction, the strain with the *cslA* plasmid exhibited significantly improved colonial growth compared to the empty-plasmid control, likely due to leaky expression from the *tipA* promoter (Fig. 4A) (Tong *et al*., 2019). Unexpectedly, *cslA* induction did not restore the overbranching phenotype associated with *divIVA* overexpression, indicating that increased CslA only partially suppressed the DivIVA overexpression phenotype (Fig. 4B, quantification Fig. S3). Plausibly, its contribution to viability may involve the enhancement of the protective layer at the cell tip.

**Figure 4.**
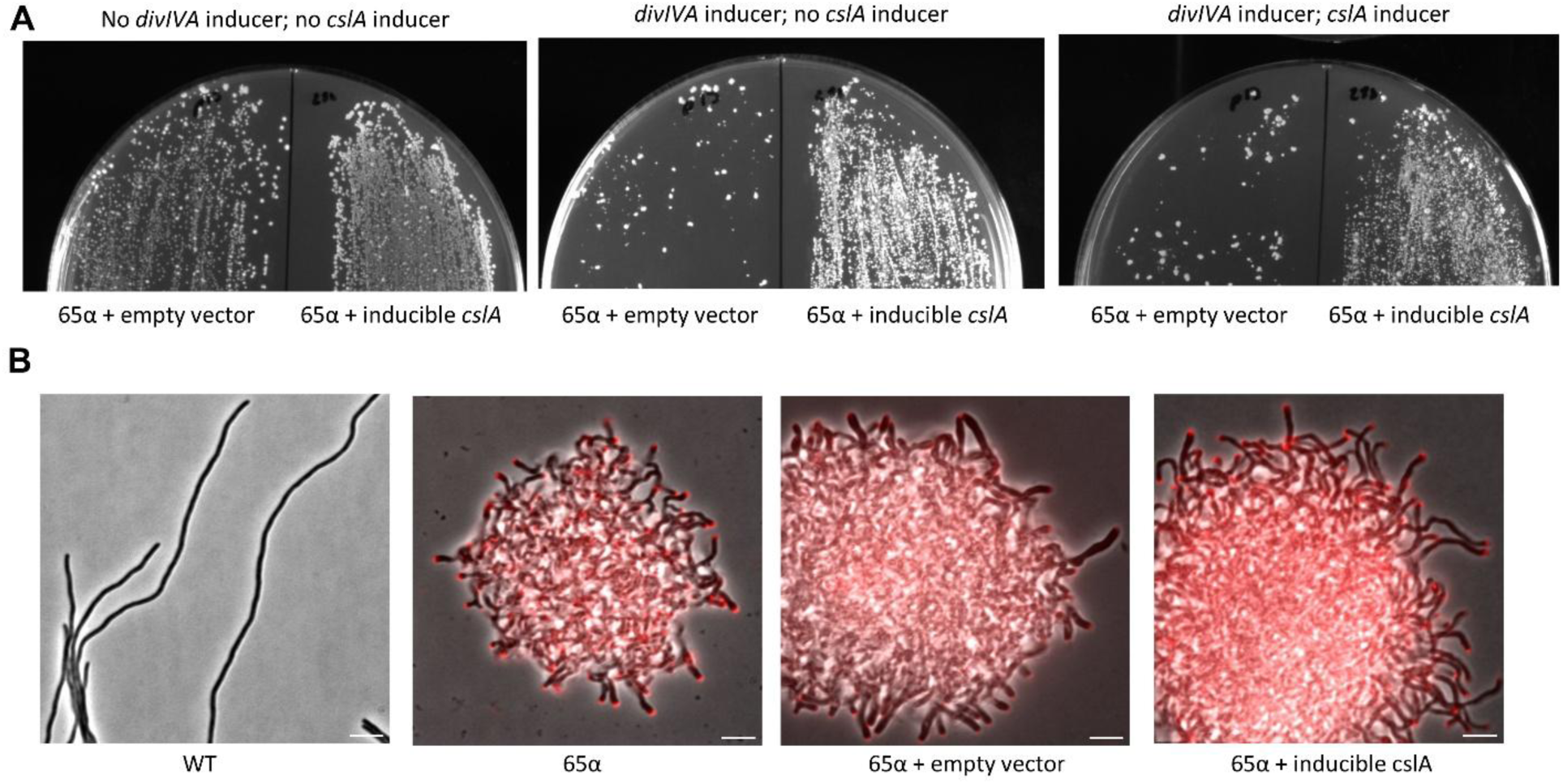
Overexpression of *cslA* rescues growth under conditions of DivIVA overproduction (A) Growth of 65α attBϕC31::pIJ6902 (empty plasmid) and 65α attBϕC31::pBK288 (*cslA* under TipA inducible promoter) grown on 5% TSB plates in the presence or absence of an inducer for DivIVA or CslA. (B) Phenotypes of *S. albus* J1074 strains (WT, 65α, 65α attBϕC31::pIJ6902 [empty plasmid], and attBϕC31::pBK288 expressing *cslA* under the TipA-inducible promoter) grown in 5% TSA medium in the presence of anhydrotetracycline (100 ng mL⁻¹). DivIVA–mScarlet production was visualised by fluorescence microscopy and is shown as a red signal overlaid on phase-contrast images. Fluorescence background and intensity thresholds were adjusted identically across all samples to allow for direct comparison.

## Discussion

To identify the factors required for tip growth and cell wall integrity, we performed a suppressor screen based on the lethal overexpression of *divIVA*, a key polarity determinant. Under these conditions, spontaneous suppressor mutants arise at low frequencies, and whole-genome- sequencing revealed recurrent mutations in the Ser/Thr kinase gene *pknB*. PknB is a conserved Ser/Thr kinase found across actinobacteria, where it has been implicated in the regulation of cell division, cell wall synthesis, and stress responses. Targeted deletion of *pknB* in the DivIVA overexpression background confirmed that the loss of this kinase is sufficient to suppress lethality (Fig. 4), indicating that PknB activity becomes problematic when DivIVA-driven tip formation is excessive.

**Figure 4.**
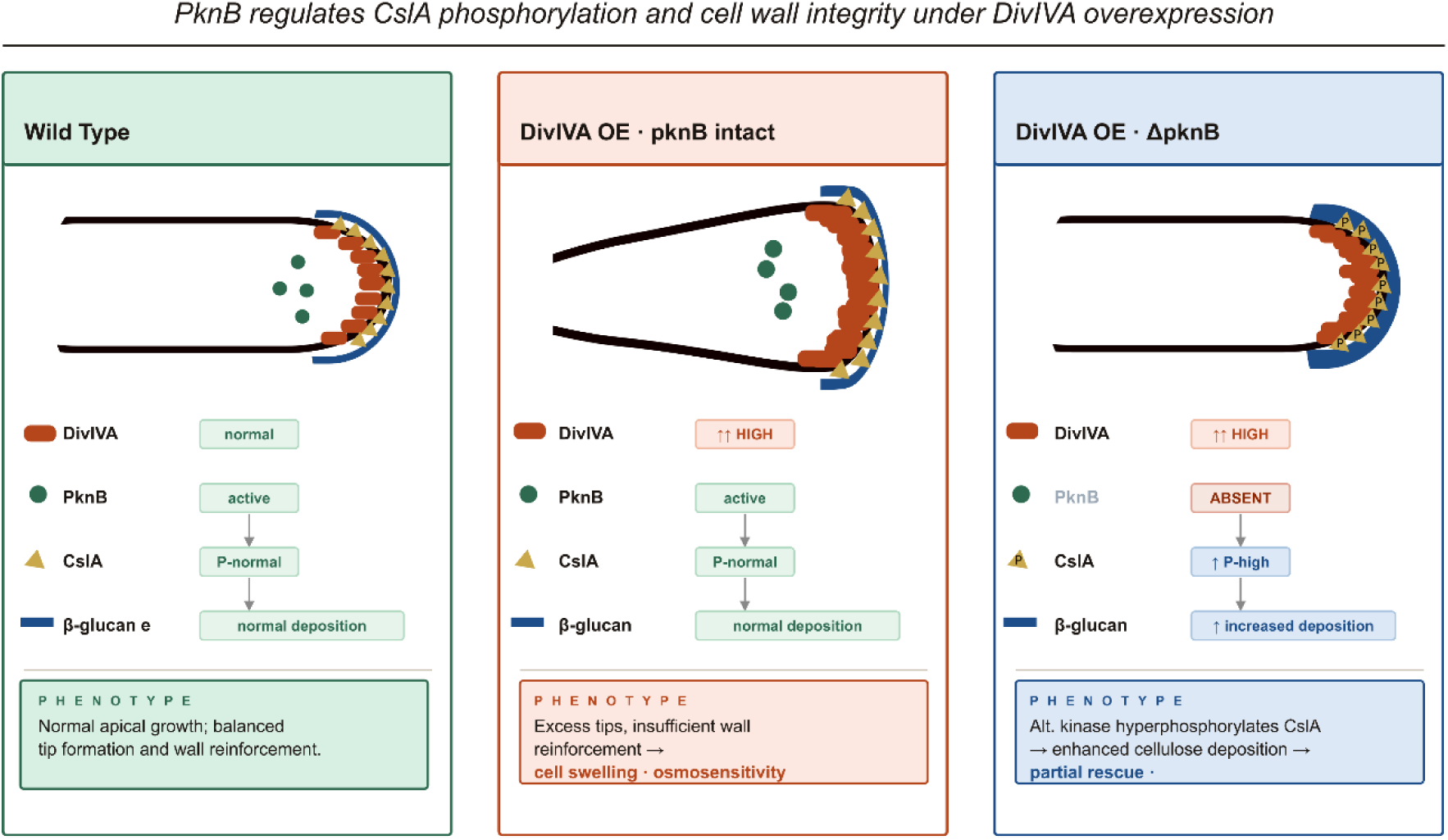
Proposed model of how phosphorylation regulates cell wall stability in *Streptomyce*s, including the proposed roles of the kinase PknB and the cellulose-like synthase CslA.

Because PknB has been implicated in diverse cellular processes, we used global phosphoproteomics to define its downstream targets. This analysis revealed that cellulose synthase CslA was hyperphosphorylated in the absence of PknB. To determine the functional relevance of this modification, we generated both a *cslA* deletion mutant and a phosphoablative T17A variant. Substitution of T17 with alanine abolished CslA activity, as shown by quantitative microscopy and increased lysozyme sensitivity, demonstrating that phosphorylation at this residue is essential for its proper function. Conversely, *cslA* overexpression partially rescued the growth defect of the DivIVA overproduction strain. However, ectopic overexpression of *cslA* did not restore normal growth and the overbranching was still detected. Therefore, additional PknB-dependent targets must contribute to the altered physiology observed in Δ*pknB* mutants. However, these findings indicate that CslA is necessary for maintaining cell wall integrity of overbranching hyphae.

These findings identify PknB as a central regulator of cell wall homeostasis that becomes essential when tip formation is elevated by DivIVA overproduction (Fig. 4). Furthermore, they implicate CslA phosphorylation as a key downstream mechanism required to maintain cell wall integrity under these conditions. The observation that CslA is more phosphorylated in the *pknB* mutant suggests that PknB modifies activity of enzyme(s) that control phosphorylation of CslA. One plausible model is that PknB represses a phosphatase or activates a second kinase, such that an alternative kinase hyperphosphorylates CslA in the absence of PknB. In *Streptomyces*, several Ser/Thr kinases, including AfsK, and the phosphatase SppA have established roles in controlling cell polarity and cell wall–associated substrates, including DivIVA. Interestingly, SppA was hypophosphorylated in our global phosphoproteome. Since SppA is known to dephosphorylate its reduced phosphorylation in the Δ*pknB* background might be expected to result in altered DivIVA phosphorylation DivIVA (Passot *et al*., 2022). However, no corresponding changes in DivIVA phosphorylation were detected in our experimental setup, suggesting that this pathway does not play a major role under the conditions tested, or that compensatory mechanisms maintain DivIVA phosphorylation levels. Whether SppA or another as-yet-unidentified enzyme mediates CslA phosphorylation downstream of PknB remains to be determined. Thus, PknB likely acts upstream of a broader regulatory network.

Finally, the DivIVA overexpression phenotype provides a physiological context for the observed regulatory circuit. Excess DivIVA generates numerous closely spaced tips, creating multiple mechanically vulnerable sites that require rapid reinforcement. The severe growth defect and pronounced low osmolarity- sensitivity of the DivIVA overproduction strain are consistent with a weakened cell envelope that is unable to withstand turgor pressure when tip reinforcement is insufficient. In this scenario, CslA becomes particularly important, and its higher levels and correct phosphorylation of CslA are likely required to deposit cellulose-rich material at all nascent growth zones. The partial suppression of the DivIVA overexpression phenotype by *cslA* overexpression fits naturally within this model; however, the failure of CslA overexpression to restore normal growth in the Δ*pknB* background indicates that multiple PknB-dependent pathways are required to maintain cell wall homeostasis when the tip number is increased.

In summary, using *divIVA* overexpression as a perturbation of tip-associated cell wall synthesis allowed us to identify PknB as a central regulator of cell wall homeostasis, acting through a phosphorylation network that ultimately controls CslA-dependent reinforcement of newly formed tips, but also involving additional PknB-regulated factors that remain to be identified.

## Author contributions

Conceptualisation (B.K., J.E.), Funding Acquisition (B.K., J.E.), Investigation (M.D., B.K, A.W., A.G., M.T.), Project administration (B.K.), Supervision (B.K., M.T., J.E.), Visualization (B.K., M.D.), Writing – original draft (M.D., B.K.,). writing – review and editing (B.K., D.J., J.E.)

## Acknowledgments

This work was funded by the Polish National Science Centre: Sonata grant 2020/39/D/NZ1/00303 to B.K. and a Wellcome Trust Investigator grant (209500) to JE.

We further wish to express our gratitude to Dr. Xiaobo Zhong (Leiden University) for useful discussions and advice regarding β-glucan staining.

We are grateful to Prof. Klas Flärdh (Lund University) for generously providing the DivIVA antiserum.

The purchase of M-Class Acquity-Synapt XS LC-MS system was financially supported by the “Excellence Initiative – Research University” program for the University of Wroclaw.

## Data availability

The genome sequencing data that support the findings of this study are openly available in NCBI at https://www.ncbi.nlm.nih.gov/sra (accession number PRJNA1099611).

The mass spectrometry proteomics data have been deposited to the ProteomeXchange Consortium via the PRIDE (Perez-Riverol *et al*., 2022) partner repository with the dataset identifier PXD053764 and PXD071342

Reviewer access details

**Project accession:** PXD053764

**Token:** tfcX8J5LlKQ2

**Project accession:** PXD071342

**Token:** jD9RJzE8ausd

