## Supplementary Data for "Roles of PknB and CslA in cell wall morphogenesis of *Streptomyces*"

#### Supplementary tables

Table S1 *Streptomyces albus* J1074 strains used in this study.

| Strain | Genotype and characteristics | Source |
|---|---|---|
| WT | *Streptomyces albus* J1074 | Gift from Prof. Dr. Andreas Bechthold (Zaburannyi *et al.*, 2014) |
| 65α | J1074; ΦBT1::tcp830-*divIVA-mScarlet* ($Hyg^R$); *divIVA*::tcp830-*divIVA-mScarlet* (native *divIVA* replaced by tcp830-*divIVA-mScarlet* cassette via CRISPR; cured) | This study |
| Δ*pknB* | J1074 Δ*pknB* (deletion of codons V39–G657); via CRISPR; cured | This study |
| 65α Δ*pknB* | J1074 *divIVA*::tcp830-*divIVA-mScarlet* (native *divIVA* replaced by tcp830-*divIVA-mScarlet* cassette via CRISPR; CRISPR plasmid cured); Δ*pknB* (codons V39–G657 deleted via CRISPR; CRISPR plasmid cured); ΦBT1-integrated tcp830-*divIVA-mScarlet*, $Hyg^R$ | This study |
| Δ*cslA* | J1074 Δ*cslA* (deletion of codons G8–L576); via CRISPR; CRISPR plasmid cured | This study |
| *cslA*$_{(T17A)}$ | J1074 *cslA(T17A)* point mutant; via CRISPR; CRISPR plasmid cured | This study |
| 65α attBϕC31::pIJ6902 | 65α attBϕC31::pIJ6902 | This study |
| 65α attBϕC31::pBK288 | J1074 *divIVA*::tcp830-*divIVA-mScarlet*; attBϕC31::pBK288 (tipA-*cslA*) | This study |

Table S2 *Escherichia coli* strains used in this study.

| Strain | Genotype and characteristics | Source |
|---|---|---|
| DH5α | F– φ80*lac*ZΔM15 Δ(*lac*ZYA-*arg*F) U169 *rec*A1 *end*A1 *hsd*R17($r_K^-$, $m_K^+$) *pho*A *sup*E44 λ⁻ *thi*-1 *gyr*A96 *rel*A1 λ⁻ | Laboratory stock (Invitrogen) |
| ET12567/pUZ8002 | *dam, dcm, hsdS,* $Cam^R$, $Tet^R$ containing plasmid pUZ8002: *tra,* $Kan^R$, *RP*4 23 | Laboratory stock (Paget *et al.*, 1999) |

Table S3 Plasmids used in this study.

| Plasmid | Genotype and characteristics | Source |
|---|---|---|
| pIJ10257 | Hygromycin resistance gene; ermE* promoter; ΦBT1 phage integration site; Origin of transfer *oriT* from RK2 | (Hong *et al.*, 2005) |
| pBK43 | Construct containing *divIVA-mScarlet* under tcp830 promoter, ΦBT1 phage integration site, hygromycin resistance gene | This study |
| DCB2202 | Synthetic construct from Dundee Cell Products containing *mScarlet* codon optimized for *Streptomyces* | This study |
| pGEMTeasy_ _Tetris_tcp | Construct containing 'tet R under the S14 promoter with the tcp830 promoter | Gift from Prof. Eriko Takano |
| pCRISPomyces2 | *Streptomyces* expression of codon-optimized Cas9 and custom gRNA, apramycin resistance gene | Gift from Huimin Zhao (Cobb *et al.*, 2015) |
| pBK64α | Construct containing *divIVA-mScarlet* under tcp830 promoter, apramycin resistance gene + protospacer detecting *divIVA* | This study |
| pBK115 | pCRISPomyces2 + protospacer detecting *pknB* + homology arms to delete 93,5% of codons in *pknB*: Val39 – Gly657 | This study |

| | | |
|---|---|---|
| pMD7 | pCRISPomyces2 + protospacer detecting *cslA* | This study |
| pMD8 | pMD7 + homology arms to exchange one codon in *cslA*: Thr17 → Ala17 | This study |
| pMD12 | pMD7 + homology arms to delete 88,5% of codons in *cslA*: Gly8 – Leu576 | This study |
| pIJ6902 | Apramycin and Thiostrepton resistance cassettes; φC31 integration site & integrase; Thiostrepton-inducible tipA promoter | (Huang *et al.*, 2005) |
| pBK288 | pIJ6902 carrying *cslA* under tipA inducible promoter | This study |

Table S4 Oligonucleotides used for cloning in this study.

| Primer | 5' – 3' Sequence | used for plasmid |
|---|---|---|
| pIJ10257_1.FOR | TGCATAGATCTAAGCTTGGATCCTAGGTTCTCA | pBK43 |
| pIJ10257_1.REV | GGTACCAGTGAGCGTTTTTCAACCT | |
| Tetris_1.FOR | GAAAAACGCTCACTGGTACCCCAAGCTTTCAACGTCAGCCG | |
| Tetris_1.REV | AGTCGTCGGCCGACGTACGCCCAATATCTCTATCACT | |
| nDiv3.FOR | GCGTACGTCGGCCGACGACTACGTTGAGG | |
| divIVa_infusion_pBK43_rv | CACCATGCTGCCGCTGCCGCTGCCGTTGTCGTCCTCGTCGATGAGGAAGCC | |
| mScralet_infusionto_pBK43_fr | GACAACGGCAGCGGCAGCGGCAGCATGGTGAGCAAGGGTGAAGCGG | |
| mScralet_infusionto_pBK43_rev | TCCAAGCTTAGATCTATGCATCACTTGTACAGCTCGTCCATAC | |
| α_top | ACGCTCAACGTAGTCGTCGGCATT | pBK64 |
| α_bottom | AAACAATGCCGACGACTACGTTGA | |
| hom_up2kb.FOR | GCAGGTCGACTCTAGAGGCGTACGCCGAGACG | |
| hom_up2kb.REV | CTGACGTTGAAAGCTGACATCACGCTCACAGC | |
| hom_dw2kb.FOR | GCATAGATCTAAGCTCGGCGCCCTAGGGTG | |
| hom_dw2kb.REV | CCGGGGATCCTCTAGAGCTCGACAACAACGTCGACT | |
| PAM_ko_RS15060_top | ACGCTGGCTGCAGATGTGCTTGGC | pBK115 |
| PAM_ko_RS15060_bot | AAACGCCAAGCACATCTGCAGCCA | |
| left_ko_RS15060.FOR | TGCCTGCAGGTCGACTCTAGAGACCTCTCCGCGCCCC | |
| left_ko_RS15060.REV | GACCGTCGCCGGCCGCGGCCGGGAC | |
| right_ko_RS15060.FOR | GGCCGCGGCCGGCGACGGTCCGGCC | |

| | | |
|---|---|---|
| right_ko_RS15060.REV | GGTACCCGGGGATCCTCTAGAGCGAGGAACTCGGACTGG | |
| oMD58 | ACGCCGTCACAGACGACCCAGCTG | pMD7 |
| oMD59 | AAACCAGCTGGGTCGTCTGTGACG | |
| oMD102 | GGAGCTGCGTGGCCTGGGAGGGGTCGGATTGGCTCCGCCC | pMD8 |
| oMD103 | CCTCCCAGGCCACGCAGCTCCGCACGGTGTCGGCCCGCACGG | |
| oMD104 | TACGGTTCCTGGCCTCTAGATGGTACTGGGCGGTGCCGAAACGG | |
| oMD105 | GCCGGGCGTTTTTTATCTAGAGGTGCGGGCCGGGGCGCGAACGGATCG | |
| oMD75 | GCCGGGCGTTTTTTATCTAGAGGGCGGCCACCGTGCG | pMD12 |
| oMD76 | GTGATCATACCCGTCGGGCTCGAC | |
| oMD77 | CGACGGGTATGATCACGGCCGCGC | |
| oMD78 | TACGGTTCCTGGCCTCTAGACGCGATGGAGCGCTG | |
| oBK391 | GTCAGAGAAGGGAGCGGACATATGACGTCGAGCCCGAC | pBK288 |
| oBK392 | GCTCGGTACCCGGGGATCCTCTAGATCATTTCTTACGTCCCCCGAGG | |

### Supplementary figures

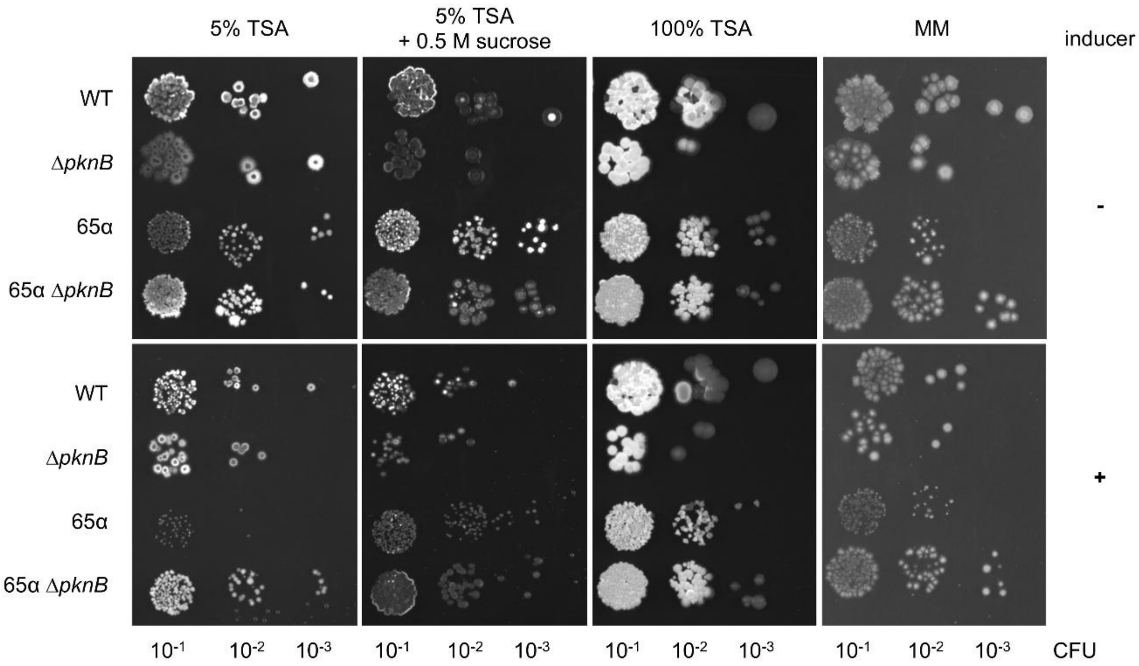

Figure S1 Survival assay of S. albus J1074.
*S. albus* J1074. strains grown on 5% TSA, 5% TSA supplemented with 0.5M sucrose, TSA or Minimal Medium (Kieser T et al., 2000) in presence or absence of the inducer - ATc 500 ng/ml.

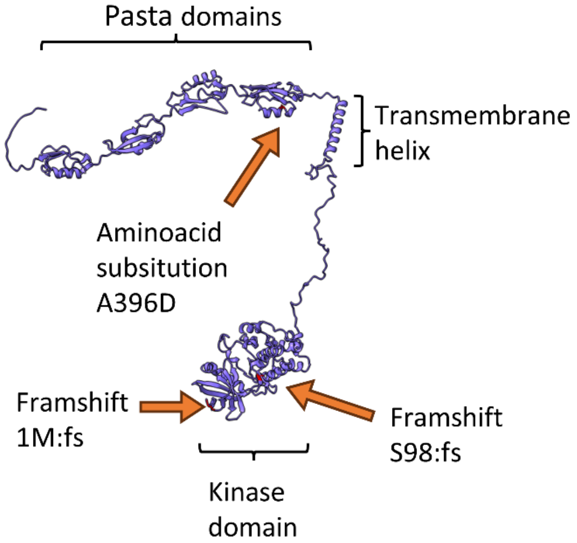

Figure S2 Alphafold model of PknB with the mutations in the suppressor screen labelled

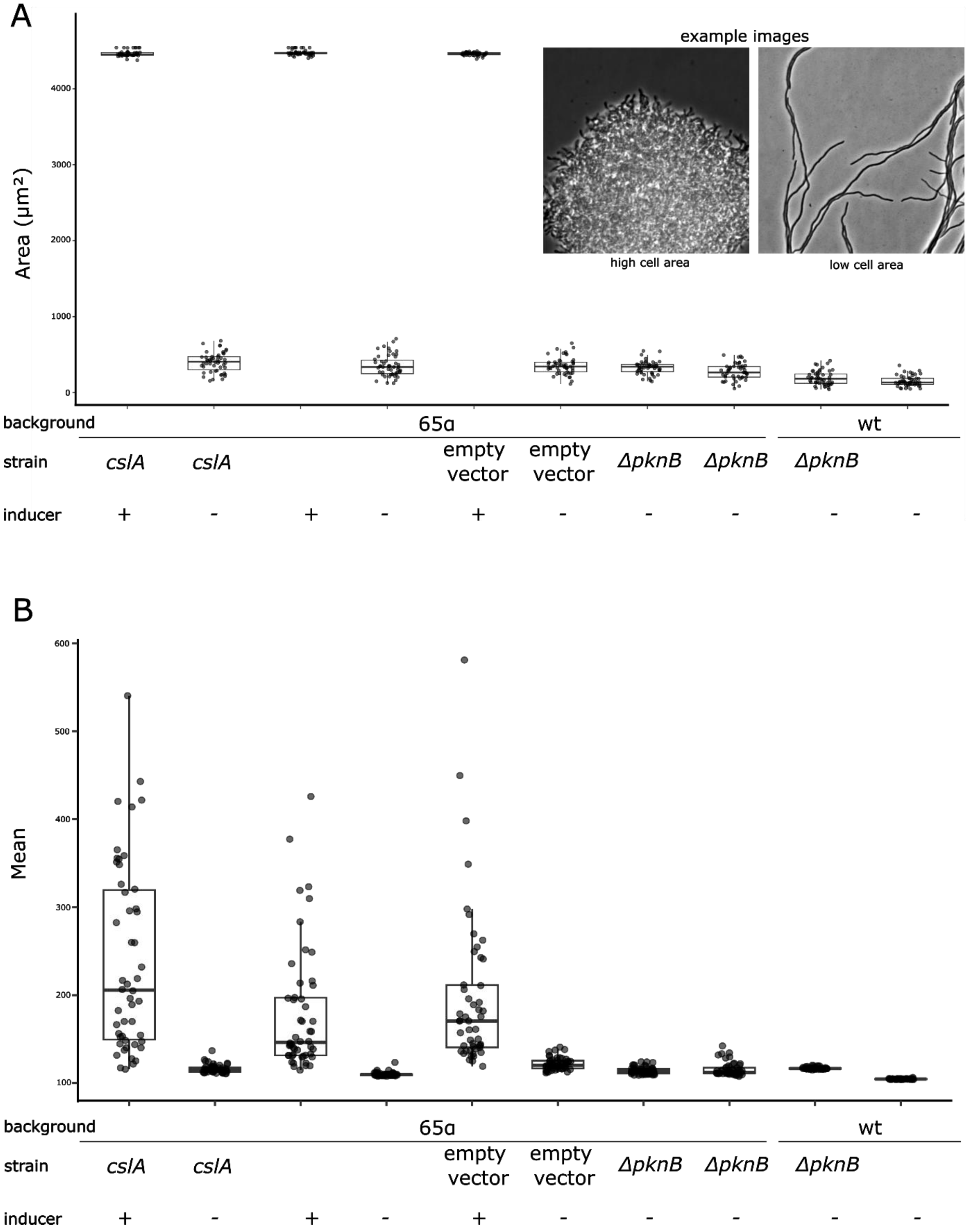

Figure S3. Quantification of microcolonies and single-cell fluorescence.

**(A)** Quantification of the microcolony area. Hyperbranching leads to larger microcolonies, which is reflected in the increased area measurements. Boxplots show the median (centre line), interquartile range (box), and 1.5× interquartile range (whiskers) of the microcolony area for each condition. Individual data points represent single microcolonies and are overlaid with jittered points. Example images of the microcolonies are shown as insets. **(B)** Quantification of

DivIVA-mScarlet fluorescence intensity in individual hyphae. Boxplots show the median, interquartile range, and 1.5× interquartile range of the mean fluorescence values for each strain. Individual data points represent single cells and are overlaid as jittered points.

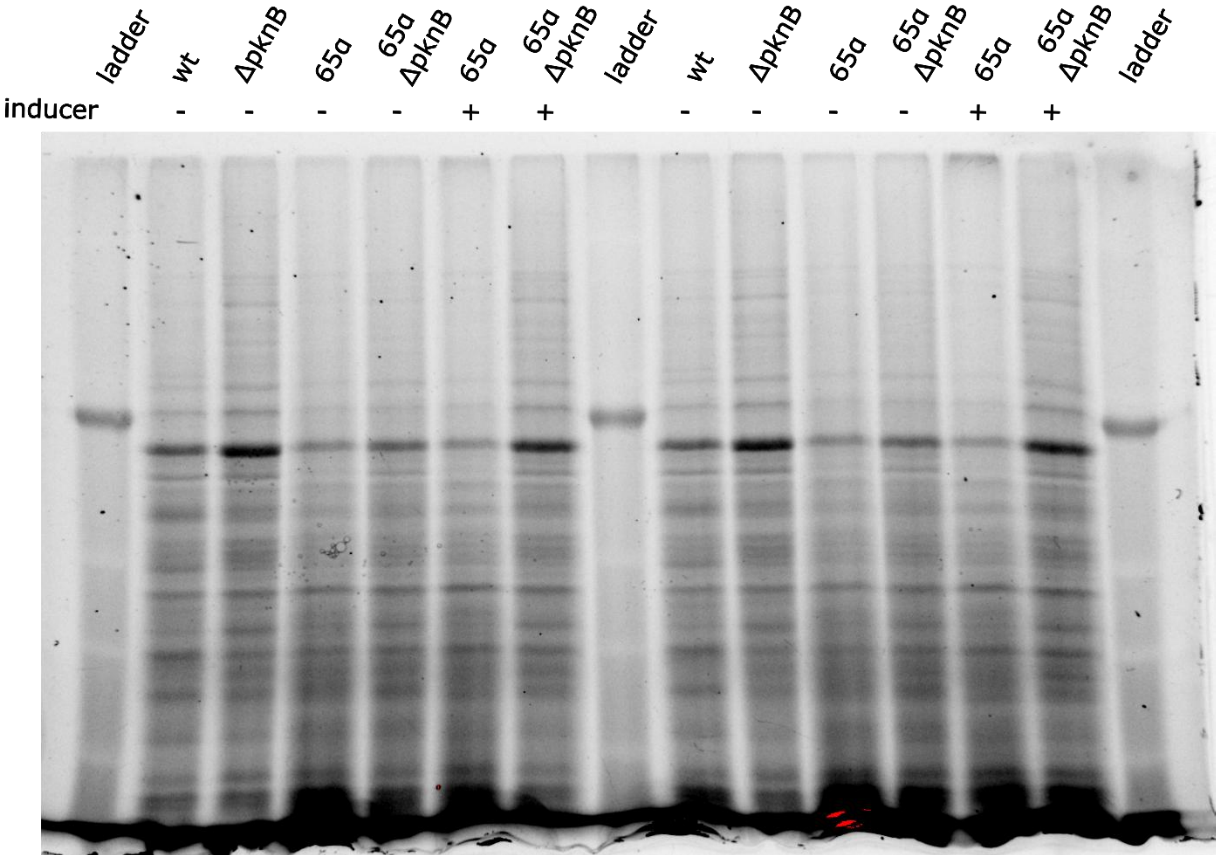

Figure S4 Stain free loading gel control.
Lanes correspond to the Western Blot results shown in Figure. 2D.

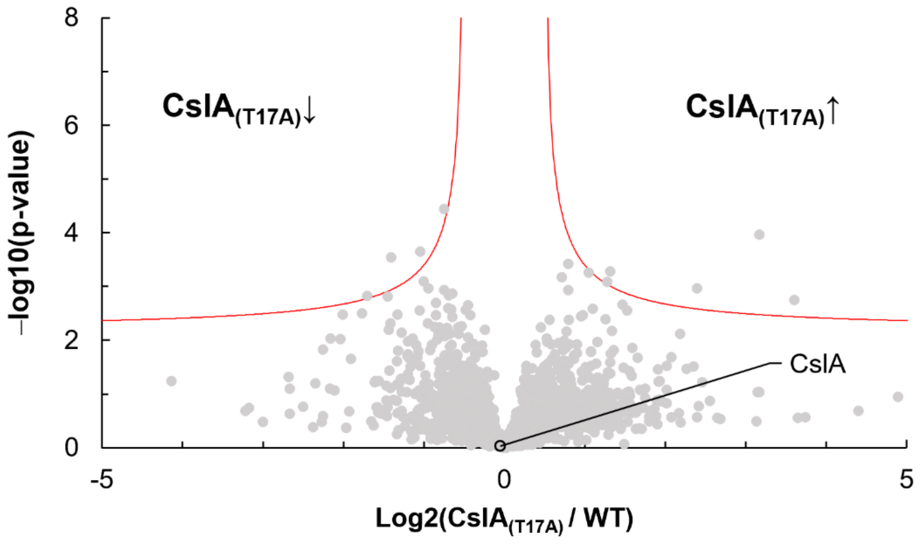

Figure S5 Volcano plot of differential protein expression in *S. albus* J1074 versus cslA$_{(T17A)}$. Each point represents a single protein that was quantified using label-free proteomics. The x-axis shows the log2 fold change (FC) in abundance between *cslA*(T17A) and *S. albus* J1074 strains, while the y-axis shows the $-\log_{10}$ of the p-value. The t-test significance criteria used for volcano analysis were an FDR of 0.05 and an S0 factor of 0.1, yielding a function for the dataset where 0.06 was the highest significant q-value and 1.67 was the lowest significant fold change. Proteins significantly upregulated or downregulated are represented by points above the red curve on the right and left sides of the plot, respectively. CslA position is indicated — CslA was quantified based on three uniquely identified peptides.

### Supplementary methods

Comprehensive Details of Plasmid Design and Assembly

**pBK43**

The vector pIJ10257 was linearised by PCR using the primer pIJ10257_1.FOR and pIJ10257_1.REV to remove the ermE* promoter. The linearised vector was then used for infusion assembly with three fragments. The first fragment contained 'tet R under the S14 promoter with the tcp830 promoter using Tetris_1.FOR and Tetris_1.REV. The second fragment comprised DivIVA from *S. albus* J1074 (primers: nDiv3.FOR and divIVa_infusion_pBK43_rv), whereas the third fragment contained *mScarlet* codon optimised for *Streptomyces* (primers: mScralet_infusionto_pBK43_fr and mScralet_infusionto_pBK43_rev). Due to primer design, the stop codon of *divIVA* was removed, and a linker encoding six amino acids consisting of alternating glycines and serines was inserted. The fully assembled construct was excised from the pIJ10257 backbone using HindIII.

**pBK64α**

First, the two protospacers (α_top and α_bottom) were annealed and inserted into pCRIPomyces-2 using Golden Gate cloning. The homology region intended for replacement was initially constructed in a pUC19 vector to facilitate easier cloning. Two 2kb fragments, located upstream and downstream of *divIVA* were generated via PCR. The following primers were used for the upstream fragment: hom_up2kb.FOR and hom_up2kb.REV. For the downstream fragment, hom_dw2kb.FOR and hom_dw2kb.REV were used. Subsequently, the fragment containing *divIVA-mScarlet* under a tetracycline-inducible promoter was excised from pBK43 using HindIII restriction enzyme. The full 6.8kb fragment was then excised from the pUC19 holding vector using XbaI and ligated into the pCRIPomyces-2 vector containing the protospacer.

**pBK115**

Using a similar approach, the two protospacers (BK_PAM_ko_RS15060_top and BK_PAM_ko_RS15060_bot) were annealed and inserted into pCRIPomyces-2 via Golden Gate cloning. The homology region intended for replacement was initially constructed in a pUC19 holding vector to facilitate easier cloning using (primers: left_ko_RS15060.FOR and left_ko_RS15060.REV; right_ko_RS15060.FOR and right_ko_RS15060.REV). The full 4kb fragment was then excised from the pUC19 holding vector using XbaI and ligated into the pCRIPomyces-2 vector containing the protospacer.

**pMD7**

Two protospacers (oMD58 and oMD59) were annealed and inserted into pCRIPomyces-2 using Golden Gate cloning.

**pMD8**

The vector pMD7 was linearized by digestion with XbaI. The homology region intended for replacement was generated via PCR. The following primers were used for the upstream fragment: oMD102 and oMD105. For the downstream fragment, oMD103 and oMD104 were used. Cloning was performed using the HiFi DNA Assembly (NEB).

**pMD12**

The vector pMD7 was linearized by digestion with XbaI. The homology region intended for replacement was generated via PCR. The following primers were used for the upstream fragment: oMD75 and oMD76. The downstream fragment was amplified using oMD77 and oMD78. Cloning was performed using the HiFi DNA Assembly (NEB).

**pBK288**

Vector pIJ6902 was linearised using the restriction enzymes NdeI and XbaI. The *clsA* gene was amplified from genomic DNA using primers oBK391 and oBK392 and subsequently inserted into the vector via sequence- and ligation- Independent Cloning (SLIC) (Li and Elledge, 2012).

Comprehensive strain construction.

**65α – DivIVA Overexpression Strain**

To generate the DivIVA overexpression strain 65α, the plasmid pBK43 was integrated into the chromosomal attB site. This construct carried *divIVA–mScarlet* under the control of the tcp830 promoter, the ΦBT1 phage integration system, and a hygromycin resistance marker. Subsequently, the CRISPR plasmid pBK64α was introduced via protoplast transformation. This plasmid contained *divIVA–mScarlet* under the tcp830 promoter, an apramycin resistance gene, and a protospacer targeting the promoter sequence of the native divIVA. The CRISPR plasmid was cleared using a temperature-sensitive origin of replication at 42 °C.

The successful construction of strain 65α was verified using Illumina whole-genome sequencing.

***Streptomyces albus* wt Δ*pknB* and 65α Δ*pknB***

*pknB* deletion strains were generated by conjugating plasmid pBK115, a pCRISPomyces2-based construct carrying a protospacer targeting *pknB* and flanking homology arms designed to remove 93.5% of the coding sequence (codons Val39–Gly657). Following CRISPR- mediated genome editing, the CRISPR plasmid was cured by incubation at 42 °C. Successful deletion of *pknB* in both the wild-type- and 65α backgrounds was verified by Sanger sequencing of 65α Δ*pknB*.

***Streptomyces albus* Δ*cslA***

The *cslA* deletion strain was generated using the CRISPR plasmid pMD12 (J1074 ΔcslA), which carries a protospacer targeting *cslA* and flanking homology arms designed to remove 88.5% of the coding sequence (codons, Gly8–Leu576). Genome editing was performed using the pCRISPomyces system, and the CRISPR plasmid was subsequently cured. The successful deletion of *cslA* was confirmed by sequencing.

***Streptomyces albus* cslA$_{T17A}$ (Thr17→Ala17 substitution)**

A point mutation- variant of *cslA* was generated using the CRISPR plasmid pMD12, which carried a protospacer targeting *cslA* and homology arms designed to introduce a single codon change (Thr17→Ala17). Genome editing was performed using the pCRISPomyces system, and the CRISPR plasmid was subsequently cured. The intended point mutation was confirmed by sequencing.

**65α attBϕC31::pIJ6902**

An empty φC31-integrating plasmid carrying apramycin and thiostrepton resistance cassettes, the φC31 integration site and integrase, and the thiostrepton-inducible- *tipA* promoter was introduced into the 65α background by conjugation. Integration at the φC31 attB site was confirmed via antibiotic selection and sequencing.

**65α attBϕC31::pBK288**

The plasmid pBK288, a pIJ6902-derived construct carrying cslA under the control of the thiostrepton-inducible tipA promoter, was integrated into the φC31 attB site of the 65α background. Integration was confirmed by antibiotic selection and sequencing of the target region.

Detailed mass spectrometry protocol

| Sample name | Label for experiment | Sample name | Label for experiment |
|---|---|---|---|
| WT_100%TSB_1/4 | WT_100_Percent_1 | ΔpknB_100%TSB_1/4 | P_100_Percent_1 |
| WT_100%TSB_2/4 | WT_100_Percent_2 | ΔpknB_100%TSB_2/4 | P_100_Percent_2 |
| WT_100%TSB_3/4 | WT_100_Percent_3 | ΔpknB_100%TSB_3/4 | P_100_Percent_3 |
| WT_100%TSB_4/4 | WT_100_Percent_4 | ΔpknB_100%TSB_4/4 | P_100_Percent_4 |

Samples were received in lysis buffer containing 8M Urea with Protease inhibitor cocktail 2 from Sigma, as recommended by the manufacturer). These samples were used for protein quantification using the BCA method, and 500 µg of protein was taken from each sample for in-solution digestion as follows:

| Samples | for 500 µg | (50 mM TEAB)100 µL | TCEP (0.5 M) 20 min RT [µL] | IAA (0.5 M) 20 min RT in dark [µL] | 50mM TEAB (10X dilution) [µL] | Trypsin (1:20) Overnight at 37°C [µg] |
|---|---|---|---|---|---|---|
| 1/4_100%_WT | 65.9 | 34.1 | 2 | 5 | 900 | 25 |
| 2/4_100%_WT | 60 | 40 | 2 | 5 | 900 | 25 |
| 3/4_100%_WT | 62.6 | 37.4 | 2 | 5 | 900 | 25 |
| 4/4_100%_WT | 68.4 | 31.6 | 2 | 5 | 900 | 25 |
| 1/4_100%_P | 62.4 | 37.6 | 2 | 5 | 900 | 25 |
| 2/4_100%_P | 45.9 | 54.1 | 2 | 5 | 900 | 25 |
| 3/4_100%_P | 58.1 | 41.9 | 2 | 5 | 900 | 25 |
| 4/4_100%_P | 51.2 | 48.8 | 2 | 5 | 900 | 25 |

Peptide clean-up for Phosphopeptide enrichment and LC-MS injection

Peptides were acidified with 10% TFA after in-solution digestion to bring the pH to <3. The samples were diluted with 3% acetonitrile and 0.1% TFA. C18 spin columns were used for sample cleanup. Briefly, the column was activated with 100% methanol (MeOH), followed by equilibration with 3% acetonitrile in 0.1% TFA. The sample was bound to the equilibrated column, followed by five washes with 3% acetonitrile in 0.1% TFA. The bound peptides were then eluted with 60% acetonitrile in 0.1% TFA, and the samples were freeze-dried in a lyophiliser. The dried samples were stored at -80°C until further use.

Phosphopeptide Enrichment

Phosphopeptide enrichment was performed using 495 µg of peptides for each replicate. In-house TiO2 based enrichment method was used. Briefly, the peptides were reconstituted in 250mM lactic acid/3% TFA/70% can. TiO2 slurry was used to prepare in-house enrichment micro-columns. The microcolumns were equilibrated with 3% TFA/70% ACN before binding the peptides to the column. The sample was then washed to remove non-phosphorylated peptides. Phosphopeptides were then eluted using 1% NH4OH in a fresh tube, dried immediately in a freeze dryer, and stored at -80°C.

LC/MS Injection

The phosphopeptides were reconstituted in 5µL of 0.1% formic acid and vortexed, sonicated, and centrifuged at 14000xg before injecting into the Mass Spectrometer, where 2 µL was injected into the mass spectrometer. Peptides were analysed by nanoflow-LC-MS/MS using a timsTOF Pro2 (Bruker) coupled to a nanoElute liquid chromatography device (Bruker). Samples were injected on a 100 µm ID × 5mm trap (Thermo Trap Cartridge 5mm) and separated on a 75 µm × 15 cm nano LC column (BrukerFIFTEEN part# 184262). All solvents used were HPLC or LC-MS Grade (Fluka™). Peptides were loaded onto the trap for 2 min at a constant pressure of 217 bar using 0.1% FA, in Water. The separation column was conditioned using 100% Buffer A (0.1% FA, in Water) at 800bar constant for 3 min and the separation was performed on a linear gradient from 2 to 25 % Buffer B (0.1% FA in Acetonitrile) over 35 minutes at 300 nL/min. The column was then washed with 90% Buffer B for 5 min and equilibrated for 10 min with 100% Buffer A in preparation for the next analysis. Full MS scans were acquired from m/z 100 to m/z 1700 at a resolution of 38000 in DDA-PASEF mode. The Ion mobility(1/K0) was measured at a range between 0.78-1.6 V.s/cm$^2$ where both the Ramp time and Accumulation time for the TIMS device was set to 100ms. The duty cycle was 100% at a ramp rate of 9.43 Hz. The same settings were used for the total proteomic experiment, where each sample was injected at a concentration of 100ng/µL.

<u>Data Analysis</u>
Peptide Identification was carried out with MaxQuant (version 1.6.10.43). MS/MS spectra were searched against the database provided earlier ("Proteins for J1074_NCBI") using a false discovery rate of 1%. The maximum missed cleavage was set to 2 using Trypsin/P enzyme. Carbamidomethylation (C) was set as a fixed modification, and acetylation (Protein N term), oxidation (M), and Phospho (STY) were set as variable modifications. The data were further analysed using a combination of R and Perseus (Tyanova *et al.*, 2016).

<u>Comments</u>
The samples were acquired for label-free quantification using a timsTOF Pro mass spectrometer. Individual two-sample t-tests for the differential expression analysis were performed for each sample in both total and phosphoproteomic experiments.
The significantly differentially overexpressed sites are marked in orange and under-expressed sites are marked in blue in all comparisons when a two-sample t-test was performed.

<u>LFQ (label-free quantitation) proteomics analysis of *S. albus* J1074 vs cslA$_{(T17A)}$ strains</u>
Samples were received in 50 mM Tris pH 7.5, 8M urea buffer. Total protein (30 µg) from each sample was denatured for 10 min at 65°C in the presence of 5 mM DTT. Samples were then diluted to 2M urea concentration with 25 mM Tris pH 7.5, 2M urea, 5 mM DTT, and 400 ng of trypsin was added for overnight digestion at 37°C. On the next day the solution was acidified and desalted using the STAGE tip procedure (Rappsilber *et al.*, 2002). The obtained peptide pellet was reconstituted in 50µL of a water solution containing 3% acetonitrile (ACN) and 0.1% formic acid (FA).

LC-MS was performed using an M-Class Acquity UPLC connected to a Synapt XS HDMS. Mobile phase A consisted of H2O + 0.1% FA, and mobile phase B consisted of ACN + 0.1% FA. For sample separation a 120 min linear gradient of 8–35%B was applied on an HSS C18 75µm × 250 mm analytical column at a flow rate of 300nL/min and column temperature set to 45°C. A 5-minute sample trapping step was performed prior to the analytical column. Six samples in total (three independent biological replicates of each strain) were run with a 5µL injection of each. MS data were collected in ion mobility DIA (HDMSE) at a scan rate of 0.6 s in the 50–2000 m/z range and Resolution Analyser Mode (~45000 FHWM at 785.84 m/z). The source conditions were fine-tuned and maintained constant. For MS2, a collision energy ramp of 27–47V was set on the instrument's transfer cell. A leucine-enkephalin solution was acquired in-parallel in the (Rappsilber *et al.*, 2002)reference function, and mass correction was applied post-acquisition.

Raw processing was performed using Progenesis QiP v4.2.7. QiP's autovalues were unchanged for the deconvolution, and the auto-optimisation of low- and high-energy ion thresholds was chosen. Signal intensity between samples was normalised according to the "Normalise to All proteins" option (robust standard deviation of each sample ion intensity estimate followed by a scalar multiplication of all ions based on that estimate). Deconvoluted data were searched using Ion Accounting against an *S. albus* protein sequence databank (CP004370) to which porcine trypsin and human keratins (UniProt entries) were appended. The set search parameters were: peptide mass tolerance: 10 ppm; fragment mass tolerance: 20 ppm; min. fragments/peptide: 1; min. fragments/protein: 3; min. peptides/protein: 1; max. protein mass: 1 MDa; digest reagent: trypsin; max. Missed cleavages: 2; variable modification: oxidation of the methionine. The FDR was adjusted to ≤1% at the peptide and protein levels. Protein grouping was performed. Peptide identifications were curated to contain only peptides with correlation coefficients ≥0.3. The intensity sum of non-conflicting peptides was used for protein quantification. For statistical analysis, protein intensity data were uploaded into Perseus (Tyanova *et al.*, 2016). The values were log2 transformed, and only proteins with at least one unique peptide and three valid values in at least one sample group were considered further, yielding a total of 1931 quantifiable proteins. Any remaining missing values (~0.25% of all values) were imputed based on the normal distribution. Reproducibility within the sample groups was ensured using PCA and Pearson's correlation tests. To identify the changed proteins, a both-sided t-test was used with significance criteria set to: q-value (permutation-based): 0.05; S0 parameter: 0.1. Detailed analysis data are available in File S4, as well as in the repository files indicated in the Data Availability section.

#### Supplementary files

File S1 Full list of mutations from suppressor screen

File S2 Mass spectrometry phosphoproteomics data

File S3 Mass spectrometry proteomics data

File S4 Mass spectrometry analysis (CslA analysis)
